# Influence of Rule and Reward-based Strategies on Inferences of Serial Order by Monkeys

**DOI:** 10.1101/2021.09.16.459819

**Authors:** Allain-Thibeault Ferhat, Greg Jensen, Herbert S. Terrace, Vincent P. Ferrera

## Abstract

Knowledge of transitive relationships between items can contribute to learning the order of a set of stimuli from pairwise comparisons. However, cognitive mechanisms of transitive inferences based on rank order remain unclear, as are contributions of reward magnitude and rule-based inference. To explore these issues, we created a conflict between rule- and reward-based learning during a serial ordering task. Rhesus macaques learned two lists, each containing five stimuli, that were trained exclusively with adjacent pairs. Selection of the higher-ranked item resulted in rewards. “Small reward” lists yielded 2 drops of fluid reward, while “large reward” lists yielded 5 drops. Following training of adjacent pairs, monkeys were tested on novels pairs. One item was selected from each list, such that a ranking rule could conflict with preferences for large rewards. Differences in associated reward magnitude had a strong influence on accuracy, but we also observed a symbolic distance effect. That provided evidence of a rule-based influence on decisions. Reaction time comparisons suggested a conflict between rule and reward-based processes. We conclude that performance reflects the contributions of two strategies, and that a model-based strategy is employed in the face of a strong countervailing reward incentive.

---

When making choices, individuals often rely on rankings of available options. Ranking may be based on prior experience or on the application of general rules. To predict the outcome of a baseball game between two teams that have never faced one another, one might consider how they fared against common opponents. Given that Team A reliably beats Team B, and Team B beats Team C, it would be reasonable to conclude that Team A would beat Team C. This is an example of a *transitive inference* (TI), the ability to infer ordinal relationships based only on implied rank without reliance on other cues (Halford et al., 2010; Piaget, 1921). Behavior consistent with TI has been observed in a wide variety of vertebrate speciesincluding rhesus macaques (*Macaca mulatta*) (Davis, 1992; Gillan, 1981; Lazareva et al., 2001; McGonigle and Chalmers, 1977; Treichler and Van Tilburg, 1996).

Transitivity is a feature of ordered sets, but it can also be a rule for making choices consistent with limited information. However, such rules may conflict with other heuristics, such as preferring items based on a subjective sense of their value. For example, some studies suggest that gamblers tend to bet on their home team even when the odds of winning are substantially less than even (Staněk, 2017). In the present study, we investigated how monkeys made decisions when a rule based on rank conflicts with learned reward values.

In behavioral paradigms that test TI, the experimenter assigns ranks to stimuli to create an ordered list (e.g. A>B>C>D>E). During training, subjects are presented with adjacent pairs from that list (e.g. AB, or BC), receiving a reward if they choose the item of “superior” rank in each pairing (Jensen, 2017). One clue that subjects are performing TI is a preference consistent with the ranking rule for novel “critical test pairs” in which both stimuli had equal reward rates during training (e.g. BD). Another is the presence of “symbolic distance effects” (SDEs, D’Amato and Colombo, 1990), in which performance improves as the gap between items’ ranks grows.

A variation of the TI paradigm is to train multiple lists in parallel, then to present new “derived lists” that are assembled from a mixture of training lists (Hakes et al., 1964). Monkeys tend to preserve the relative ranks of items presented in new derived combinations (Swartz et al., 1991; Chen et al., 1997; Treichler and Raghanti, 2010; Jensen et al., 2020). This suggests that they rely on linear representations of rank to evaluate otherwise ambiguous derived pairs (Merritt and Terrace, 2011; Jensen et al., 2020; Mione et al., 2020).

Although the mechanisms that result in TI remain unclear, reinforcement learning (RL) provides a framework for characterizing potential strategies. Algorithms that make choices only by associating stimuli with expected reward values are “model-free” because they do not rely on any other organizational structure (Dayan and Niv, 2008; Jensen et al., 2019b). Model-free learning therefore depends on reward history, and cannot exploit rules based on the assigned ranks of stimuli (Ferrucci et al., 2019; Minamimoto et al., 2009; Stanisor et al., 2013). Model-free strategies fail to describe behavior in many TI paradigms (Lazareva and Wasserman, 2006, 2012; Gazes et al., 2012; Vasconcelos and Monteiro, 2014; Jensen et al., 2019a).

As an alternative approach, “model-based” algorithms assume that stimuli relate to one another within a representational framework and that a subject’s choice is based on rules relating to that representation (Dayan and Niv, 2008). Model-based accounts of TI that rely on the representation of an item’s position along a continuum predict most TI behaviors, including SDEs (Kao et al., 2020; Jensen et al., 2020; Mione et al., 2020). This continuum has previously been described as a cognitive representation of a serial order (Terrace, 2010, 2012).

Jensen et al. (2019a) showed that Q-learning, a model-free algorithm, cannot solve classic TI, either when all rewards are equal or when low-ranked items are associated with larger rewards than high-ranked items. However, Q-learning succeeds when differential rewards are concordant with the ranking rule (Jensen et al., 2019a,b). This suggests that model-free RL can be a useful heuristic in some TI contexts, even though behavior in TI tasks is otherwise well-described by model-based RL (Smith and Church, 2018).

In this study, we measured the contributions of reward-based and rule-based components of learning and decision-making by training subjects using lists for which the magnitude of reward differed. The conflict between reward disparity and relative item rank in derived pairs allowed the dissociation of rule-based and reward-based contributions to preference.

## Methods

### Subjects

Subjects were three adult male rhesus macaques (*Macaca mulatta*) identified as O, R and S and housed at Columbia University. The study was carried out in accordance with the guidelines provided by Guide for the Care and Use of Laboratory Animal of the National Institute of Health (NIH). The Institutional Animal Care and Use Committee (IACUC) at Columbia University approved the study.

All subjects had previous experience with serial learning procedures, including transitive inference tasks. Subjects earned rewards that were delivered in units of “drops-” with each drop having a volume of approximately 0.20mL. Typical performance during a session earned between 100mL and 250mL, whereas perfect performance could earn up to 300mL. Subjects also received a ration of biscuits (provided before the session) and fruit (provided after the session).

### Apparatus

Subjects performed the task using a touchscreen that was linked to a computer. The subjects sat in a specially designed primate chair while performing the task. The touchscreen (model 1939L, Elo Touch Solutions, California) had a 19” display (1280×1024 resolution at 60 Hz) with a resistive touchscreen interface to record responses. All tasks were programmed in Matlab (2019, Mathworks, Natick MA) using the Psychophysics Toolbox (Brainard, 1997; Kleiner et al., 2007; Pelli, 1997). To deliver fluid rewards, the computer was connected to a solenoid valve through an Arduino Uno interface. Fluid was delivered through a stainless-steel tube attached to the primate chair.

### Stimuli and Task

Stimuli were presented on the touchscreen. Subjects responded by touching them when they appeared. The stimuli were photographic images drawn from a large set of royalty-free stock images. They were grouped into randomly drawn lists of five images, with an ordering pre-defined by the experimenter. We denote the item positions using the labels ABCDE, with A being the highest rank and E the lowest rank. No indication of stimulus rank was presented to subjects beyond that implied by correct/incorrect feedback. Additionally, stimulus lists were inspected prior to the experiment to ensure that stimulus features were not predictive of list order. The images were presented in pairs, one image on the left and one on the right side of the display. Only two stimuli were presented on each trial. Stimulus arrangement on screen was counterbalanced over trials to avoid left/right bias.

Figure 1A presents the experimental design. Each trial began with the presentation of a solid blue square (100 × 100 pixels) at the center of the screen to focus the attention of the subject and to provide a consistent target for their hand. After the subject touched the square, two stimuli (200 × 200 pixels) were presented on either side of the screen. These stimuli always differed in rank. Touching the stimulus with the higher rank was “correct” and resulted in a fluid reward and the presentation of a green check as visual feedback. Touching the stimulus with the lower rank was “incorrect” and resulted in presentation of a red X, which was followed by a time-out lasting 2s. During the time-out, the screen was blank. For example, when presented with the pair AB, a reward could only be earned by touching the stimulus A.

**Figure 1.**
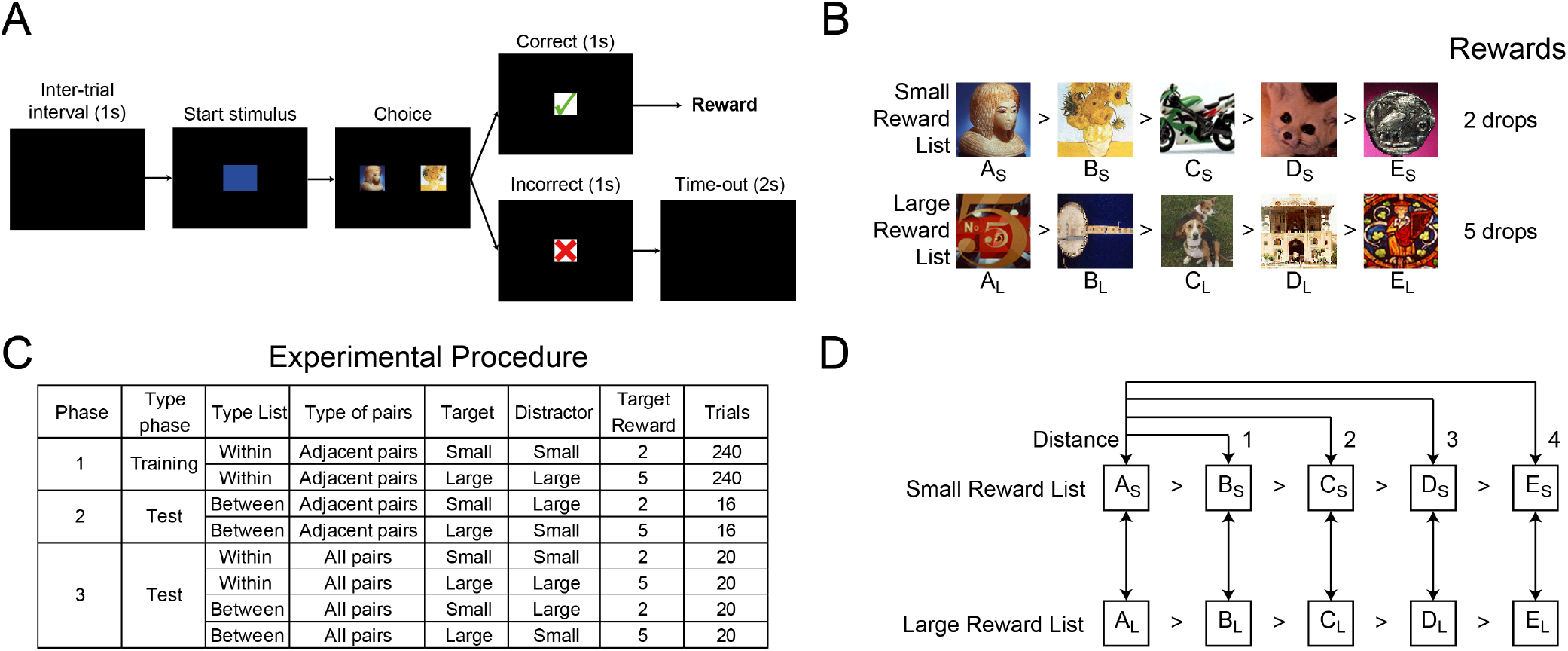
Description of the experimental procedure. **(A)** Representation of trial procedure. The trial started with the presentation of a blue square after a one-second inter-trial followed by the presentation of pairs from within or between lists. If the subject touches the higher-ranked item, a reward is delivered according to the type of list associated. Otherwise, a time out of 1-3 seconds occurred. **(B)** Example of the five-item list annotated small list and large reward list and the reward associated, consisted of randomly picked stock image. **(C)** A detailed description of the type of pairs presented during each of the three phases. All types of pairs appearing during the same phase were intermixed with one another. **(D)** Schematic description of the interaction between all items according to the distance and the possibility of switch items between the small reward list and large reward list.

Two 5-item lists (with ranks denoted by ABCDE, e.g. Figure 1B) were used in each experimental session. The 10 images used in these two lists had not been seen in any prior session and were not re-used in subsequent sessions. To study the effect of reward magnitude on learning, correct responses on one list were rewarded with 5 drops of fluid (the “large reward” list); on the other, with 2 drops (the “small reward” list). We denote which list a stimulus belongs to with a subscript, such that AS indicates the highest-ranked item in the small reward list, whereas EL indicates the lowest-ranked item in the large reward list. As a control, there was also an “equal reward” condition, during which subjects still learned two lists but all correct responses yielded rewards of 2 drops of fluid.

Each subject completed 25 sessions per condition, one session per day. Each session comprised a training phase, followed by two different testing phases. During training, subjects were presented trials with adjacent pairs where the two items were from the same list (“within list” pairs). This yielded eight pairs: A_S_B_S_, B_S_C_S_, C_S_D_S_, D_S_E_S_, A_L_B_L_, B_L_C_L_, C_L_D_L_, and D_L_E_L_. While each trial presented only items from one list, the two lists were randomly interleaved across trials. Trials were organized into blocks, during which each pair was presented twice: once with the target (higher ranked item) on the left, and once on the right. The sequence of presentations within a block was randomized. Training continued in this fashion for 30 blocks, or 480 trials.

During the first testing phase, a new set of adjacent pairs was assembled by taking one item from each list (“between list” pairs). This yielded eight new pairs: A_S_B_L_, B_S_C_L_, C_S_D_L_, D_S_E_L_, A_L_B_S_, B_L_C_S_, C_L_D_S_, and D_L_E_S_. When evaluating choices made for these derived pairs, the amount of reward was determined by which list the correct item was drawn from. If the subject was presented with the pair A_S_B_L_, choosing A_S_ yielded a reward of 2 drops and choosing B_L_ yielded a time-out. However, if the subject was presented with the pair A_L_B_S_, choosing A_L_ yielded a reward of 5 drops and choosing B_S_ yielded a time-out. These eight pairs were also arranged into blocks of trials, such that presentations within each block randomly permuted the order of presentation. Each pair appeared twice per block, once with the target on the left, and once with the target on the right. This first testing phase consisted of two blocks, for a total of 32 trials.

During the second testing phase, all possible pairs were presented, including both within-list pairings such as B_S_D_S_ and between-list pairings such as C_S_E_L_. Since 10 pairs can be drawn from 5 items, there were 10 within-list small reward pairs, 10 within-list large reward pairs, 10 between-list pairs with a small reward target, and 10 between-list pairs with a large reward target. The presentation order of each pair was randomized, with all pair types intermixed and with counterbalanced target positions. Thus, the second testing phase comprised a total of 80 trials. Figure 1C summarizes these trial counts and compositions. Figure 1D gives examples of each of the four types of pairings that can occur in this experiment overall: adjacent within-list pairs (symbolic distance of 1, e.g. A_S_B_S_, seen during training), adjacent between-list pairs (symbolic position of 1, e.g. A_L_B_S_, seen during the first testing phase), and all-pairs combinations that included both within-list pairs and between-list pairs (e.g. A_S_E_S_ and C_S_E_L_, seen during the second testing phase).

If the subject failed to initiate a trial by pressing the start stimulus or initiated and then waited more than 4s to make a choice, the trial was not counted toward the requirement for completing that phase and was presented again. For the unequal reward condition, within-list adjacent-pairs presentation, an average of 562.98 trials were attempted, relative to the 480 completed trials necessary to complete the phase trials (O: 541.36, R: 664.56, S: 483.04); for between-list adjacent-pairs presentations, 46.21 trials were attempted of the 32 required (O: 59.64, R: 42.12, S: 36.88); for all-pairs presentations, 100.53 trials were attempted of the 80 required (O: 106.84, R: 105.12, S: 89.64). For the equal reward condition, within-list adjacent-pairs presentations, 562.98.24 trials were attempted of the 480 required (O: 542.72, R: 761.68, S: 481.92); for between-list adjacent-pairs presentations, 46.21 trials were attempted of the 32 required (O: 42.72, R: 40.24, S: 37.04); for all-pairs presentations, 100.53 trials were attempted of the 80 required (O: 97.48, R: 112.04, S: 89.16).

### Analysis

Each subject’s performance, quantified as proportion of correct responses, was modeled using logistic regression with trial number, symbolic distance, and reward size as explanatory variables. These variables are related to outcomes using the following logistic transformation:

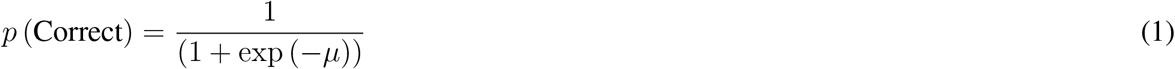

Here, *μ* represents a linear combination of terms, which varies with the phase of the task being modeled. Unless otherwise specified, individual predictors that contribute to *μ* use mean-centered values in order to avoid creating strong covariance with intercept terms. Consequently, in all models described below, the intercept (denoted by *β*_0_) corresponds to a ‘typical’ case, and each predictor corresponds to some deviation from that typical case.

Since our chief interest was to assess the contributions of both rule-based and reward-based influences on behavior, we relied on a regression model to tease apart these contributions. One of the important rule-based effects we expected to observe was the “symbolic distance effect” (SDE) (D’ Amato and Colombo, 1990). When a subject displays an SDE, their response accuracy, or reaction times, or both change as a function of the relative rank of the items. For example, given a list ABCDE, a positive effect of distance would result in BE (a distance of 3) having higher accuracy than BD (a distance of 2). SDEs are frequently observed in the empirical literature, and are consistent with the view that TI relies on a cognitive representation of list order (Terrace, 2012; Jensen, 2017). We operationalized *D* as our measure of symbolic distance in regressions that include pairs of multiple distances. Although the symbolic distance between items varied from 1 to 4, smaller symbolic distances were more numerous than large ones. Because of this, *D* was centered by subtracting 2 from every case, as that was the mean distance of all possible pairs. For example, the value of *D* for the pair AB was – 1 = (1 – 2), whereas the value of *D* for the pair AD was 1 = (3 – 2). This centering ensured that estimates of *β_D_* did not covary with other parameters any more than necessary.

Similarly, we operationalized our reward manipulation as *R*, a predictor that was a centered and dummy-coded, with *R* = 0.5 when the target stimulus belonged to the large reward list and *R* = −0.5 when the target belonged to the small reward list.

During training, every trial used adjacent pairs, which have a symbolic distance of 1. Since there was no variation in distance among these pairs, training performance was modeled based on the learning rate in terms of trials *t* and the reward amount *R*, as well as the interaction of these two variables. We categorized training data as “early” (first third of trials), and “late” (last third of trials), to allow the model to best fit to data. This yielded the following linear equation during training:

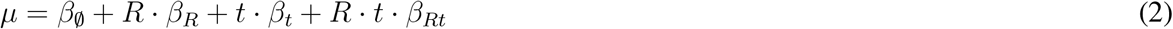

Here, *β*_0_ is an intercept term that describes overall response accuracy at the start of training. As noted above, *β_R_* describes any differential response accuracy observed at the start of training. *β_t_* acts as a learning rate, describing how response accuracy changes over time, and *β_Rt_* describes the differential learning rate between the small reward list and the large reward list.

The first testing phase (between-list adjacent-pairs) consisted of too few trials for a learning rate *β_t_* to be estimated. With this in mind, we used the following linear combination:

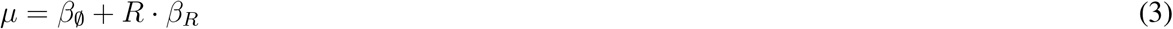

Since trials are omitted as a predictor, *β*_0_ and *β_R_* are more appropriately interpreted as descriptions of overall performance during the adjacent-pair between-list testing phase. As above, *R* was coded as the reward magnitude associated with the target, with a value of 0.5 when the target belonged to the large reward list and −0.5 when the target belonged to the small reward list.

During the final testing phase, trials were again omitted as an explanatory variable, and performance was predicted in terms of *R*, the reward size associated with the target. Additionally, the symbolic distance *D* was also used as a predictor, resulting in the following linear combination:

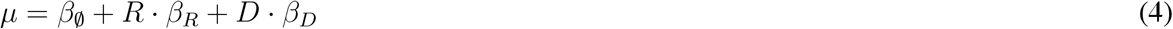

As noted above, *D* was a centered predictor, and as such estimates of *β_D_* did not covary with either estimates of *β_R_* or of *β*_0_.

During the equal reward condition, the parameters of reward *β_R_* and *β_Rt_* were removed from the equations described above because R did not vary in that condition. Additionally, we treated performance for list 1 and list 2 as interchangeable with respect to targets for equal reward condition data. As such, A_1_B_1_ and A_2_B_2_ were both simply treated as within-list pairs, whereas A_1_B_2_ and A_2_B_1_ were both treated as between-list pairs.

Parameters were estimated using multi-level models, with each regression parameter estimated for each subject, as well as at the population level. These models were implemented in the Stan programming language (Stan Development Team, 2019; Carpenter et al., 2017) and scripted using the R language (R Foundation for Statistical Computing, Vienna).

## Results

To study the impact of reward disparity on learning and decision-making, we first trained subjects on two ordered lists of 5 items each, where each trial presented a pair of items drawn from one list or the other (Figure 1). Each daily session comprised an initial training phase followed immediately by the testing phases. During training, only within-list adjacent pairs were presented, with either small or large reward size associated with each pair for unequal reward condition (Figure 1C). The different possible pairs were then presented during the following test phase (Figure 1C). Each daily session used 10 new items on which subjects had not been trained previously.

During training, performance reached 75% correct responses for both small and large reward lists (Figure 2A). Only the CD pairs showed lower performance compare to other adjacent pairs, a pattern consistent with past reports in the literature (Jensen et al. 2019a). Thus, when subjects learned two lists in the same session, reward magnitude did not seem to have an effect on overall performance at the end of training, as long as both items presented on each trial belonged to the same list.

**Figure 2.**
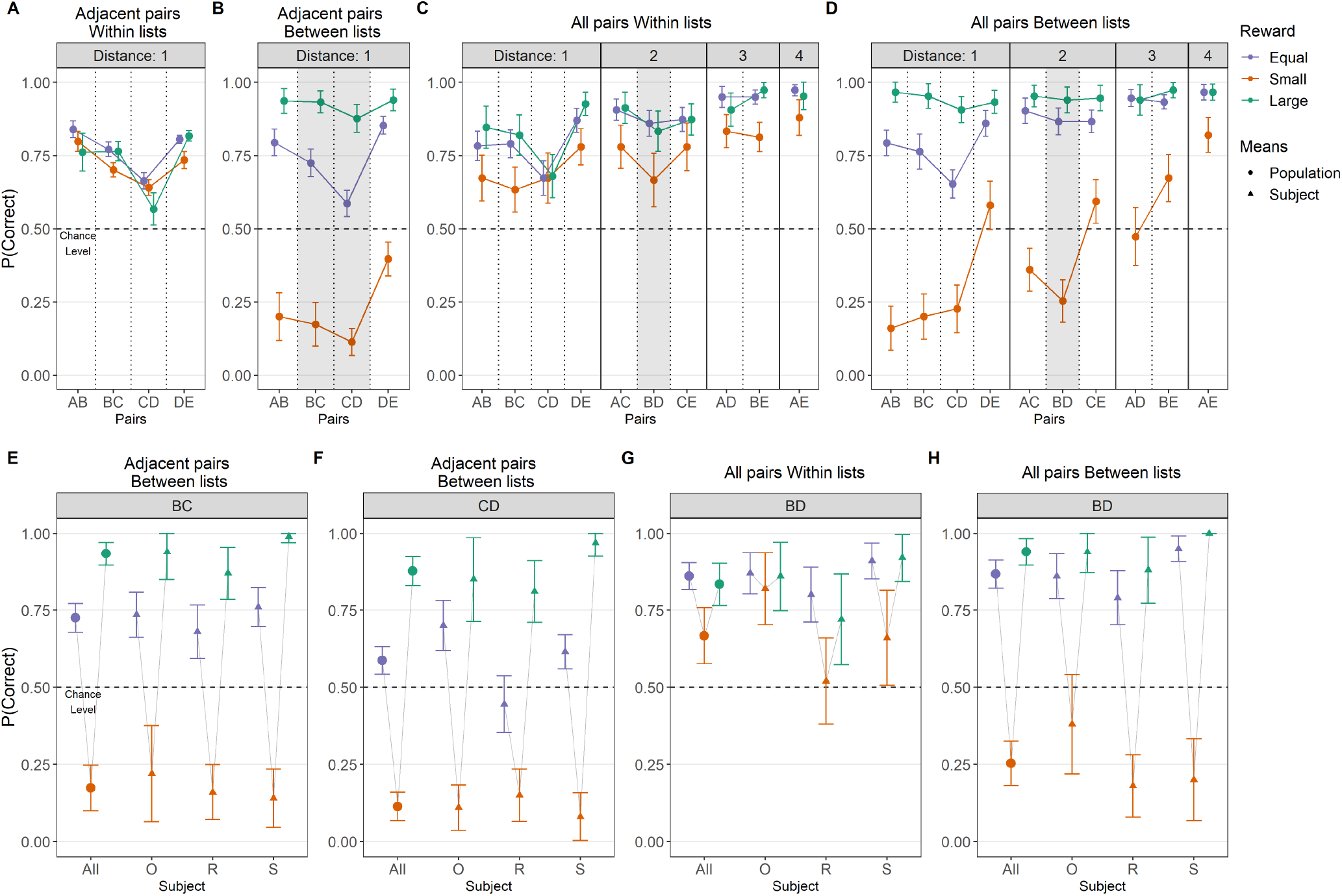
Observed average response accuracy of each type of pairs (A-D) and critical pairs (E-H) for each reward manipulation. Points and error bars depict the average and 95% confidence interval for observed performance for each pair. Performance **(A)** during training, adjacent pairs within list presentation, **(B)** during adjacent pairs between list presentation, **(C)** during all pairs within list presentation, **(D)** and during all pairs between list presentation. **(E-H)** Performance of critical pairs during equal, the small reward condition and the large reward condition for the population and each subject: BC and CD pairs during between-list adjacent-pairs presentations **(E & F)** and BD during within-list **(G)** and between-list all-pairs presentations **(H)**. Pairs are annotated accordingly to type of reward, Equal condition (purple), small rewarded target (orange) and large rewarded target (green).

We then observed the performance for three types of test pairs: “adjacent between-list pairs”, “all within-list pairs”, and “all between-list pairs”. For test pairs comprising adjacent items drawn from different lists with the same reward magnitude, subjects were able to perform well above chance (Figure 2B, purple line), consistent with previous studies (Terrace, 2010). This establishes that subjects were able to make correct between-list comparisons when reward was not a confound. We have previously referred to this ability as “positional inference” (Jensen et al., 2021).

However, when a target item from the small reward list was presented with a distractor item from the large reward list, adjacent pair performance was far below chance levels (Figure 2B, orange points). In other words, non-human primates tended to choose the item that had been associated with the larger reward during training, even though this went against the rule of choosing the item with higher rank. Such choices were not rewarded during testing, but rather resulted in a time-out.

Conversely, in adjacent pairs where the target was from the large reward list and the distractor from the small reward list, performance was not only above chance, but also better than performance in the equal reward condition (Figure 2B, green points). All conditions displayed a similar pattern of higher performance for terminal pairs (AB and DE) and lower performance for CD pairs (Figure 2B), despite their different baseline accuracies.

Performance for all combinations of items from both lists (all collected simultaneously during the final testing phase) is here partitioned into within-list pairs (Figure 2C) and between-list pairs (Figure 2D). This testing phase included pairs with symbolic distances of 1 (adjacent pairs) to 4. In all cases, a symbolic distance effect (SDE) was evident, with performance generally increasing as a function of symbolic distance.

Furthermore, for the equal reward condition, within- and between-list performance was nearly identical, providing further evidence of positional inference. Performance for within-list pairs was also similar when comparing the equal reward condition to the unequal pairs with large reward targets (Figure 2C). Performance for pairs with small reward targets, however, was consistently lowest of the three (Figure 2C, orange line).

As in Figure 2B, performance for between-list pairs with small reward targets during all-pairs testing was below chance in many cases (Figure 2D). Despite this, subjects still displayed SDEs, and as such small reward target pairs were selected above chance when the symbolic distance became large enough. Pairs with large reward targets showed the opposite reward effect, with elevated performance relative to the equal reward condition, effectively reaching a performance ceiling.

Figures 2E–2H plot the performance for “critical test pairs”, those novel adjacent or non-adjacent pairs that omit both the first and the last items (e.g. BD or CD, but not AC or BE). Performance for pairs with large reward targets was consistently at or above 75% correct, and either equal or superior to performance in the equal reward condition. However, small reward target pairs were below chance in all between-list cases (Figures 2E, 2F & 2H), and showed a disadvantage even in within-list comparisons, despite mostly being above chance (Figure 2G).

Overall, performance strongly suggest that subjects understood the rule to “choose the higher rank,” and were able to do so both when rewards were equal and when they were concordant with expected reward magnitude (Jensen et al., 2019a). Furthermore, even when a rule-reward conflict drove performance below chance, the presence of SDEs during all-pair testing suggests that subjects still demonstrated some understanding of the list order, despite the biasing effect of the reward disparity. This selective deployment of a rule-based strategy raises the question of how the rule interacted with reward associations, particularly when the two were in conflict. To address this, we examined reaction times.

Figure 3 shows reaction times on a log scale for all pairs (Figures 3A–3D) and critical pairs (Figures 3E–3H), split for correct/incorrect responses. In general, during all phases and regardless of whether correct or incorrect, mean reaction times for the equal reward condition had the same magnitude (Figures 3A–3D, purple points). An overall SDE is also visible, with faster reaction times for large symbolic distances.

**Figure 3.**
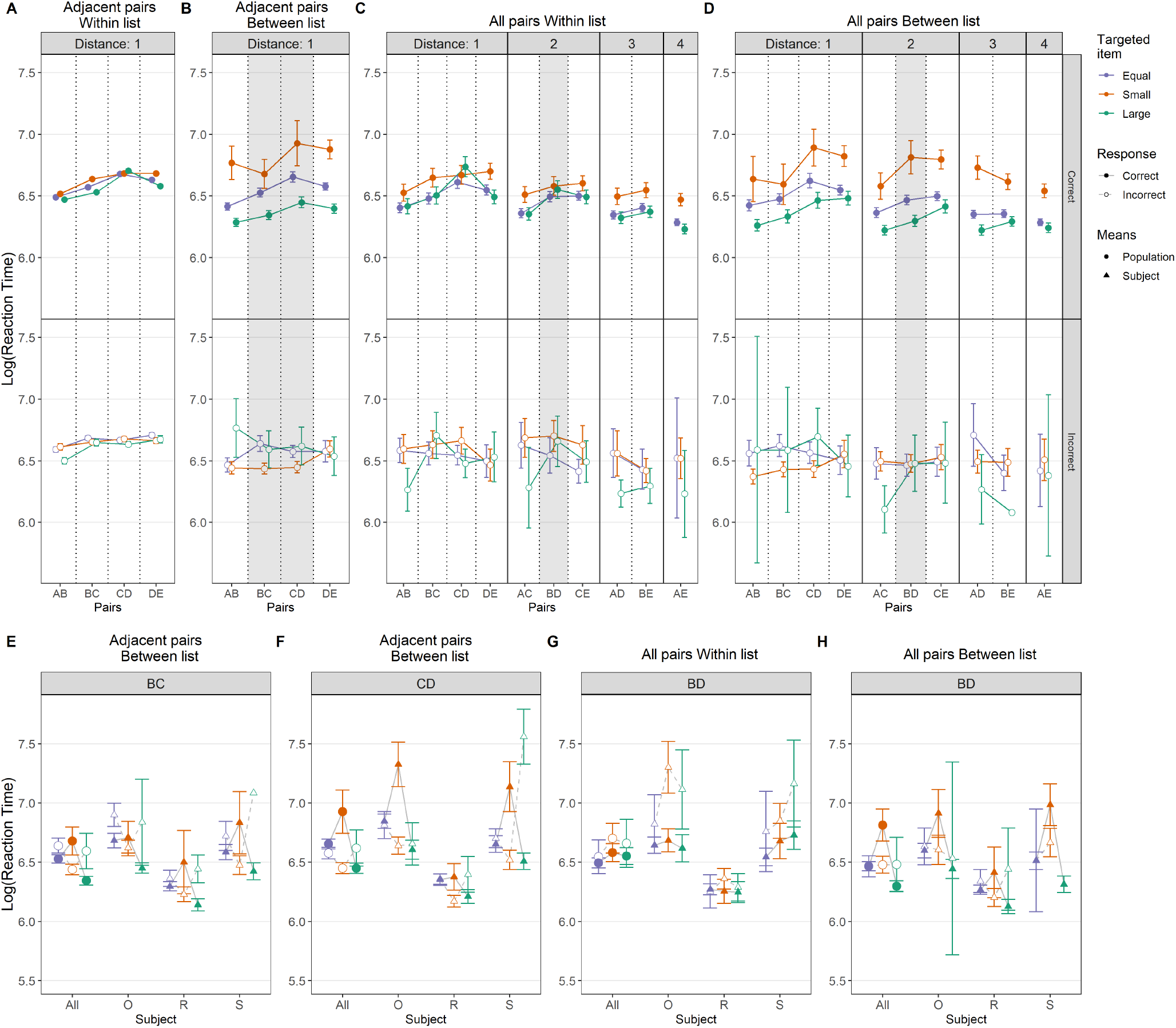
Observed average reaction time of each type of pair (A-D) and critical pairs (E-H) for each subject and each reward manipulation. Filled points correspond to reaction times for correct responses, whereas open points correspond to reaction times for incorrect responses. Points and error bars depict the average and 95% confidence interval for observed reaction time for each pair. Mean log reaction times **(A)** during training, adjacent pairs within list presentation, **(B)** during adjacent pairs between list presentation, **(C)** during all pairs within list presentation, **(D)** during all pairs between list presentation. **(E & F)** Mean log reaction times for the pairs BC and CD during between-list adjacent-pairs presentations for equal, the small reward target and the large reward target. **(G & H)** Reaction time for the pairs BD during within-list and between-list all-pairs presentations. Pairs are annotated accordingly to type of reward, Equal condition (purple), small rewarded target (orange) and large rewarded target (green).

However, the introduction of a reward disparity induces a substantial split in reaction times. Between-list pairs with small reward targets are, when selected correctly, also chosen substantially more slowly than incorrect responses (Figures 3B and 3D, filled points). Even the within-pair correct trials of the testing phase show a slowdown for small reward targets when they are paired with small reward distractors (Figure 3C, filled points). In contrast, when the target is associated with the larger reward, reaction times for correct responses were comparable to equal reward trials for within-list pairs and faster than equal reward trials among correct between-list pairs (Figures 3B & 3D, filled points).

In summary, when a small reward target was paired with a large reward distractor, correct responses were particularly slow and deliberate, whereas when a large reward target was paired with a small reward distractor, correct responses were particularly fast.

Figures 3E–3H directly contrast the correct and incorrect response reaction times for the critical pairs, BC, CD, and BD, doing so for both overall means and for individual subjects. In the equal condition, differences in reaction times between correct and incorrect responses were small or nonexistent. For all between-list means, small reward trials (orange) were slow when the response was correct, and were fast when the response was incorrect (Figures 3E, 3F, and 3H). The opposite was true for the large reward trials (green). This difference in reaction time suggests an increased difficulty and a cognitive cost to respond during decisions in which the rule and the reward were in conflict. A competition between rule and reward-based decision processes, rather than just an overall tendency to respond more quickly to items associated with larger rewards, seems to explain this result. Note that because subject S had a 100% correct response rate to large reward targets in between-list pairs (Figure 2H), we cannot estimate that subject’s reaction time for incorrect responses (Figure 3H).

To quantify the relative contributions of rule and reward-based decision processes, we fitted performance data with the regression model described in Equations 1 to 4 to each phase. This regression revealed three aspects of the learning: (a) *β*_0_, the performance at the beginning of each phase (or the phase overall when no learning rate was included) (b) *β_D_*, the impact of increased distance between pairs on performance, associated with the SDE and the rule-based decision process, and (c) *β_R_*, the impact of reward disparity on performance, a proxy of the contribution of reward-based process. Figures 4A, 4C, and 4E show model parameter estimates for the unequal reward condition in each phase of the experiment, while Figures 4B and 4D do the same for the equal reward condition. At the beginning of each session (“Adjacent Within Early”), the intercept *β*_0_ was at chance levels for each subject in each condition (unequal and equal reward, Figure 4A & 4B “Adjacent Within Early”). During late stage training and all subsequent test phases, values of *β*_0_ were consistently positive, demonstrating that subjects performed above chance overall, whether there was a reward disparity or not. Performance that is reliably above chance on trial 0 indicates that information has been learned and retained from the previous phase of the experiment.

**Figure 4.**
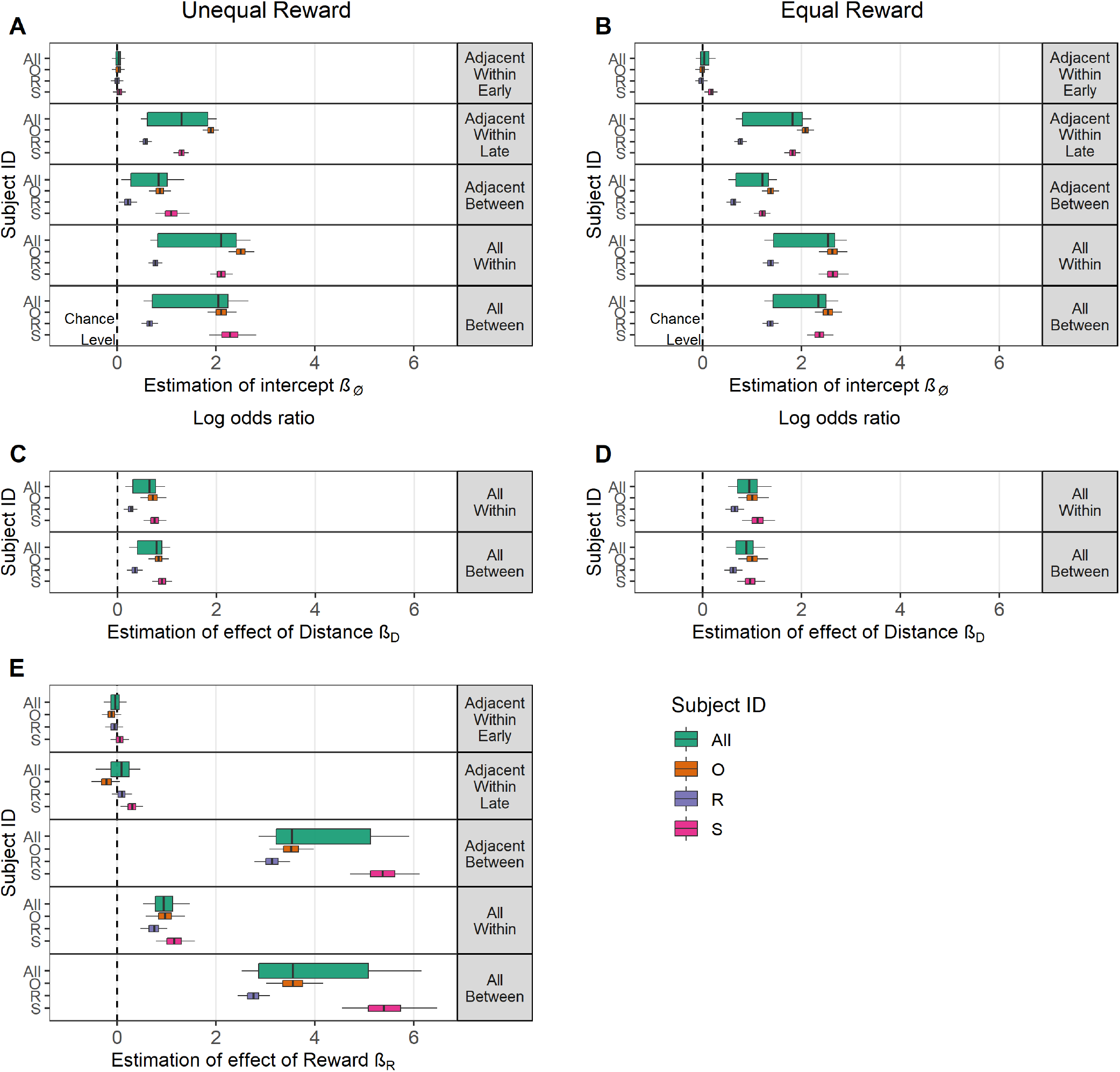
Estimated logistic regression parameters for the population (green box) and each subject (O: orange, R: purple and S: pink) for unequal (Left panels) and equal reward manipulation (Right panels). Boxes represent the 80% credible interval, and whiskers represent the 95% credible interval. **(A & B)** Estimates of the intercept *β*_0_, describing performance at trial 0 for each type of pair presentation for the unequal reward condition (Left) and equal reward condition (Right). **(C & D)** Estimates of *β_D_*, describing the impact of symbolic distance for the unequal reward (Left) and equal reward (Right). **(E)** Estimates of *β_R_*, describing the impact of reward disparity for the unequal reward condition.

The estimates of *β*_0_ for “Adjacent Between” reflect performance during between-list testing. Values above zero indicate that the representation of order learned during training transferred to testing, insofar as positional inferences among adjacent pairs are concerned. *β*_0_ was lower for the unequal rewards condition than for the equal rewards condition, suggesting that the reward disparity in some way tended to impair transfer.

The predictor *β_D_* captures the effect of symbolic distance during all-pairs testing. This effect was consistently positive and the 95% credible intervals exclude zero (Figure 4C & 4D) with performance increasing with larger symbolic distance. In the unequal reward condition, the SDE was robust for both within-list and between list pairs (Figure 4C, “All Within” vs. “All Between”). If anything, the between-list SDE was slightly greater than that for within-list transfer. The SDE was stronger in the equal reward condition (Figure 4D) than it was in the unequal reward condition (Figure 4C), but was equally strong for all test pairs belonging to each reward condition. Overall, these results show a consistent and durable effect of symbolic distance, even when differential rewards produced strong biases in responding.

Figure 4E plots estimates of the reward disparity effect *β_R_* in the unequal reward condition. *β_R_* was close to 0.0 during training (Figure 4E “Adjacent Within Early” & “Adjacent Within Late”), suggesting that subjects did equally well on the training pairs for both lists even with reward disparity. However, a dramatic reward effect appeared when between-list adjacent pairs were tested (Figure 4E, “Adjacent Between”), and this persisted throughout all-pairs testing (Figure 4E, “All Between”) indicating that subjects had a strong bias for the item drawn from the large reward list. Interestingly even the within-list pairs shown this effect during all pairs testing (Figure 4E “All Within”), reflecting higher performance on large reward pairs than small reward pairs, a result seemingly at odds with the comparable performance for both lists during the training phase. This last detail can’t be explained by a mere “reward preference,” and may signal a broader effect on motivation.

Figure 5 compares the observed empirical means of pairs to the model fits calculated using the regression parameters described above. Figures 5A–5C shows the mean probability of a correct response during blocks of training trials for each condition during both early and late training. In general, observed data were in close correspondence with the model description, although some curvature evident in the first three or four blocks of the empirical data were not quite captured by the regression model.

**Figure 5.**
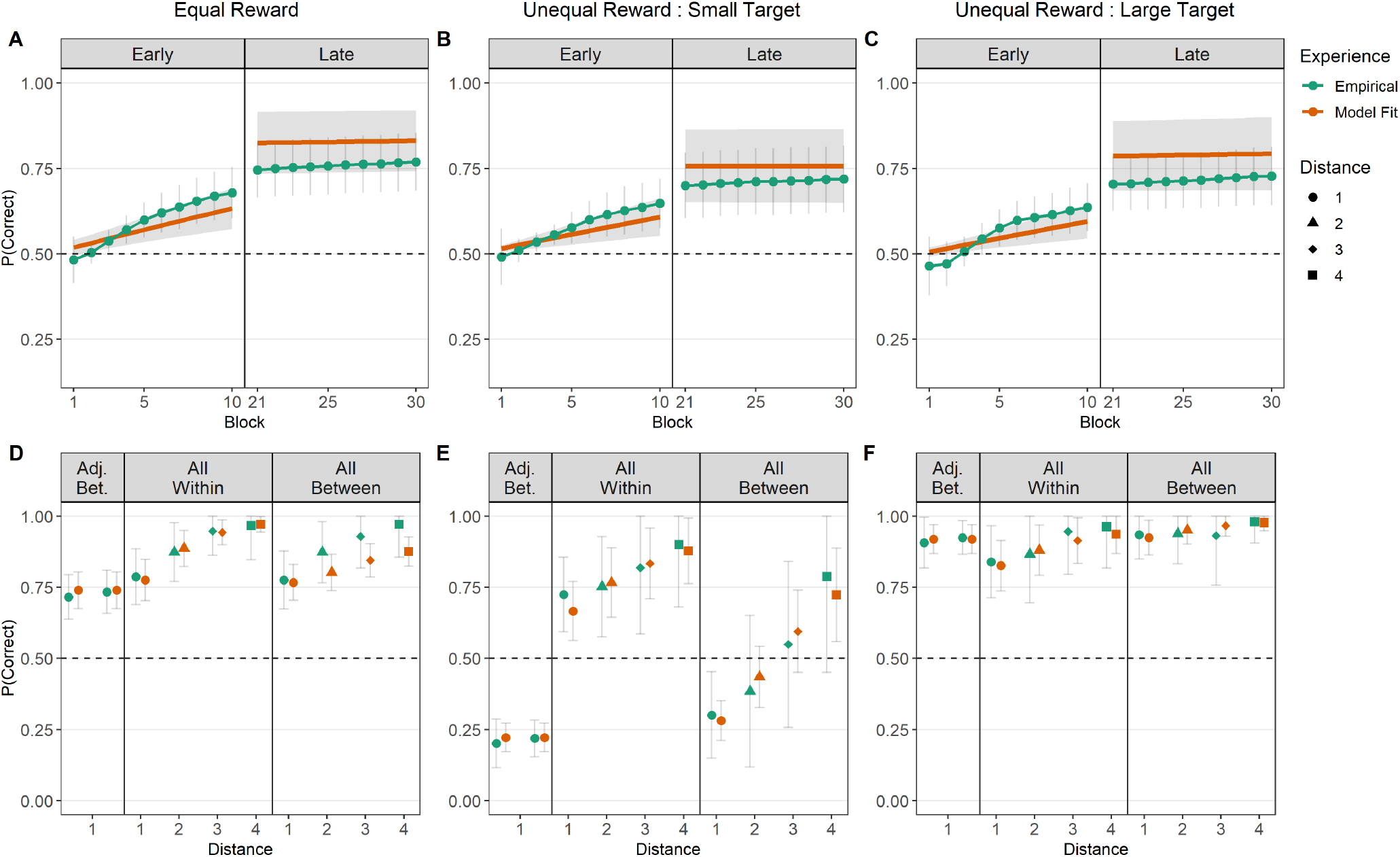
Empirical (green points) and model fit (orange lines and points) performance during training (A: equal, B: small reward, C: large reward) and testing (D: equal, E: small reward, F: large reward). Points represent average observed performance during each block, whereas lines represent best-fit model estimates. Error bars and shaded regions depict 95% credible intervals. **(A & D)** Performance during equal reward presentations. **(B & E)** Performance when the targeted item was from the small reward list during unequal reward presentations. **(C & F)** Performance when the targeted item was from the large reward list during unequal reward presentations.

A similarly good correspondence was found between empirical means and model fits of the testing phases. Above-chance responding and an evident SDE were captured in the equal reward condition (Figure 5D). Similarly, the interaction between symbolic distance and reward disparity yielded accurate recapitulation of both the interference effects when subjects were presented with small reward targets (Figure 5E) and the near-ceiling performance when subjects were presented with large-reward targets. In general, these results indicate that the parameters plotted in Figure 4 provide a good overall summary of the contributions of distance and reward effects, and as such give a suitable summary description of the data.

## Discussion

There is a long-standing controversy as to whether performance on serial learning tasks reflects the application of transitive inference, or can instead be explained by model-free reinforcement learning. We investigated the cognitive mechanisms underlying serial learning by putting reward-based and rule-based comparisons of stimuli into conflict with one another. The goal was to measure the relative influence of these two types of learning. It might have been the case that subjects always favored a rule-based strategy, even when it forced them to choose an item previously associated with a low reward. Conversely, subjects might have forsaken the application of transitivity and chosen only according to experienced reward associations.

Instead, the results revealed a complex interplay of rule- and reward-based strategies. When rewards were equal, it was clear that monkeys could apply the rule both within and across lists. However, when the stimulus was associated with a small reward, it was difficult for monkeys to choose that item when it was paired with a large-reward stimulus, even when the ordering rule that had been established during training dictated that they should do so. Such choices were suboptimal in that they resulted in no reward. However, subjects also displayed a critical hallmark of transitive inference, the symbolic distance effect (SDE), such that overcoming the biasing effect of differential rewards was easier when item ranks differed by a larger amount. Reaction times were much slower when making correct choices when these two influences were in conflict than when the influences were concordant, suggesting competition between rule, and reward-based decision strategies. The lack of a clear reaction time difference in the equal reward condition demonstrates that such large reaction time differences are not a typical feature of TI, and thus point specifically to an impact of reward disparity on the decision-making process.

Prior to introducing between-list test pairs with differential rewards, performance was broadly consistent with past reports of TI in rhesus macaques. When other factors were controlled for, subjects responded above chance overall at test and performed similarly on within- and between-list pairs in most cases, suggesting that subjects did not find derived pairs difficult in general and that they relied on positional inference to solve novel pairings. It is noteworthy that the *β*_0_ parameter was similar in both the equal and unequal reward conditions and reliably above chance following early training, suggesting that subjects displayed comparable overall learning whether or not a reward disparity was present during training (Figure 4A & 4C).

Subjects also displayed positive symbolic distance effects (SDEs), as measured by the *β_D_* parameter, in both equal and unequal reward conditions. Although the size of this effect was slightly smaller in the unequal reward condition, the effects were of a similar magnitude (Figure 4C & 4D). Thus, despite a powerful overall preference for stimuli taken from the large reward list (Figure 2B), subjects responded at greater than chance levels of accuracy to the most widely spaced pairs, as the SDE effectively overtook the reward bias for the largest symbolic distances. This result is consistent with past studies of TI, both those using standard paradigms and those using derived lists (Jensen et al., 2020; Terrace, 2012). The presence of both above-chance performance to test pairs and reliably positive SDEs point is not only evidence of TI, but, more broadly, it is evidence of a stable rule-based component of serial learning. This is in keeping with previous model-based proposals that serial learning relies on a representation of stimulus position along a continuum.

Interestingly, despite dramatic effects of reward disparity, the estimated intercept (*β*_0_) and effect size of the symbolic distance effect in our model (*β_D_*) was similar in both the unequal and equal reward conditions, with the latter only slightly larger. However, despite estimates of *β*_0_ and *β_D_* providing clear evidence of TI, subjects also displayed a very large effect of reward disparity, as measured by the *β_R_* parameter (Figure 4E). Subjects displayed a strong preference for stimuli associated with the large reward list, resulting in enhanced accuracy presented with large reward targets, but reduced (and often below-chance) accuracy when presented with large reward distractors (Figure 2B & 2C, “Adj. Bet.” and “All Between”). While these effects “average out” to above-chance performance overall, the effect of differential reward still dominates preference in most cases. Furthermore, discrepancies in reaction time between correct and incorrect responses in the unequal reward condition (Figure 3B–3C) suggest a competition between rule-based and reward-based strategies that were cognitively costly when the “target” (according to the rule) was associated with small rewards.

The effect size of the reward manipulation was much larger than that reported by Jensen et al. (2019a), who also put reward discrepancies in conflict with the ordinal rank of list items. Subjects may have had more success overcoming the reward bias in that study because, unlike the present study, they were trained on single lists with a unique reward for each stimulus. Thus, the derived list paradigm (and thus using positional inference to combine knowledge across multiple lists) may be more vulnerable to this kind of disruption than transitive inference alone. Another possible factor is that subjects in the present study received less training for each list than was the case in the 2019 experiment, where each list was trained for multiple sessions prior to transfer.

Surprisingly, the effect of reward was also positive during within-list all-pairs testing (Figure 4E), resulting in lower within-list performance for test items from the small reward list. Since within-list small reward stimuli did not have reward discrepancies with respect to one another, this must reflect some broader effect on learning induced by reward disparities across trials. For example, subjects may have exhibited differential motivation for (or paid differential attention to) stimuli from one list relative to the other. Reward magnitude has played an important role in modulating the learning rate during other tasks (Ferrucci et al., 2019; Vartak et al., 2017), and thus may do more than merely increase a response bias. Furthermore, several studies show that differential reward magnitudes impact motivation and attention (reviewed by Jovanovic and Matejevic, 2014).

Following these interpretations, the *β_R_* observed during between-list presentations could be a mixture of (a) variations in attention and motivation that influenced learning and (b) a response bias arising directly from the non-informative reward disparity itself. At a minimum, a purely reward-associative account would only predict an increased *β_R_* during between-list presentations. As such, this differential learning may be another clue as to why subsequent between-list reward biases were as large as were observed. A future study with a more direct index of either motivation or attention could clarify to what extent these are modulated by differential rewards.

Although differential rewards had a powerful influence on preference, a strictly model-free account is not sufficient to explain the behavior observed in this study. Both the consistently positive SDEs and greater-than-zero intercept terms suggest that model-based learning was active and stable during training and, to some extent, independent of reward associations. Rhesus macaques appear, on average, to favor rule-based choices even when expected rewards conflict with the rule (see also Gazes et al., 2012). We interpret our results as suggesting that the representational strategy was being applied by subjects in both conditions, and to similar degrees, independent of the confounding effect of differential rewards. However, based on our analysis of reaction time, the influence of reward on accuracy seems to be somewhat in competition with the rule-based representation, rather than reward value being fully and directly integrated into each subject’s representation of order. Rule-based and reward-based contributions having a degree of independence may also explain why such consistent distance effects (as measured by *β_D_*) were observed, even as performance on specific pairs was disrupted.

Our results make clear that a full picture of learning must incorporate reward-based associative effects, while retaining the transitive characteristic of a cognitive model. While two competing learning systems may be at play, a theory of TI should ultimately incorporate both effects, rather than treating these factors as each being evidence for mutually exclusive theories of behavior.

Our present results are also consistent with previous demonstrations that subjects rely on positional inference when evaluating between-list pairs (Jensen et al., 2020; Terrace, 2012), doing so to a similar degree both in the presence and absence of differential reward effects. Thus, we attribute the low response accuracy for between-list critical pairs during the small reward trials (Figure 2D–2G) as being due to reward bias instead of errors in positional inference. The presence of a positive SDE in between-list all-pairs presentations (Figure 2B, 4C & 4D) supports the understanding of the rules and the presence of a reward bias.

Overall, we observed that monkeys seem to use both model-free and model-based reinforcement learning to perform transitive and positional inference. The reward effect seems to interact with the representation of the lists and the rules according to the symbolic distance, generating an improvement of learning and better performance for the large reward list than is observed during classic TI performance (i.e. the equal reward condition). However, when ordinal position and expected reward are in conflict, as in between-list pairs when the targeted item is from the small reward list, the two types of learning, model-free and model-based, are also in conflict relative to both the amount of reward and the informativeness of the rule (in this case, the symbolic distance between items). It is, therefore, not surprising to observe, the presence of both clear reward biases and of differential reaction times as a function of conflict between these factors. These findings point to considerations that should be explored using TI tasks that employ derived lists. Since performance reflects the influence of both rules and reward, an important objective is to better understand the ways in which these learning systems interact in serial learning.

## ACKNOWLEDGMENTS

The authors would like to thank Dr. Fabian Munoz for his assistance and advice on collecting data and writing the manuscript, and Anna Meaney for her assistance in training animals and collecting data.

## AUTHOR CONTRIBUTIONS

ATF, GJ, HST, and VPF conceived the experiments. ATF acquired data. ATF and GJ performed the analyses. ATF, GJ, HST, and VPF wrote the paper.

